# Psilocybin-induced changes in brain network integrity and segregation correlate with plasma psilocin level and psychedelic experience

**DOI:** 10.1101/2021.02.03.429325

**Authors:** Martin K. Madsen, Dea S. Stenbæk, Albin Arvidsson, Sophia Armand, Maja R. Marstrand-Joergensen, Sys S. Johansen, Kristian Linnet, Brice Ozenne, Gitte M. Knudsen, Patrick M. Fisher

## Abstract

The emerging novel therapeutic psilocybin produces psychedelic effects via engagement of cerebral serotonergic targets by psilocin (active metabolite). The serotonin 2A receptor critically mediates these effects by altering distributed neural processes that manifest as increased entropy, reduced functional connectivity (FC) within discrete brain networks (i.e., reduced integrity) and increased FC between networks (i.e., reduced segregation). Reduced integrity of the default mode network (DMN) is proposed to play a particularly prominent role in psychedelic phenomenology, including perceived ego-dissolution. Here, we investigate the effects of a psychoactive oral dose of psilocybin (0.2-0.3 mg/kg) on plasma psilocin level (PPL), subjective drug intensity (SDI) and their association in fifteen healthy individuals. We further evaluate associations between these measures and resting-state FC, measured with functional magnetic resonance imaging, acquired over the course of five hours after psilocybin administration. We show that PPL and SDI correlate negatively with measures of network integrity (including DMN) and segregation, both spatially constrained and unconstrained. We also find that the executive control network and dorsal attention network desegregate, increasing connectivity with other networks and throughout the brain as a function of PPL and SDI. These findings provide direct evidence that psilocin critically shapes the time course and magnitude of changes in the cerebral functional architecture and subjective experience following psilocybin administration. Our findings provide novel insight into the neurobiological mechanisms underlying profound perceptual experiences evoked by this emerging transnosological therapeutic and implicate the expression of network integrity and segregation in the psychedelic experience and consciousness.

## Introduction

Current neuroscientific models informed by functional neuroimaging studies hold that multiple macroscale cerebral functional networks, through coordinated activity, shape behavior and phenomenal experience (i.e., subjective experience of contents in consciousness) (Lord et al., 2017; Margulies et al., 2016a; Smith et al., 2009). These functional networks are reliably identified using blood-oxygen-level-dependent (BOLD) resting-state functional magnetic resonance imaging (rs-fMRI) (Damoiseaux et al., 2006; Raichle, 2011; Yeo et al., 2011), an imaging technique for measuring functional connectivity (FC), which is a proxy for neural communication (Biswal et al., 1995). An axiomatic feature of these functional networks is high FC between nodes belonging to the same network (“network integrity”) and low FC between nodes belonging to different networks (“network segregation”) (Raichle, 2011). Reduced network integrity and segregation has recently been proposed to underlie the psychedelic experience after intake of psilocybin (Carhart-Harris and Friston, 2019) but neuroimaging experiments evaluating these relations are sparse (Mason et al., 2020; Roseman et al., 2014; Tagliazucchi et al., 2014).

The psychedelic compound psilocybin potently induces an altered state of consciousness and is emerging as a promising novel therapeutic, showing long-lasting beneficial effects with fast onset after a single dose both in healthy individuals (Griffiths et al., 2018, 2011; Maclean et al., 2011; Madsen et al., 2020) and in patients with depression, anxiety and addiction (Bogenschutz et al., 2015; R. L. Carhart-Harris et al., 2016; Carhart-Harris et al., 2018; Griffiths et al., 2016; Grob et al., 2011; Johnson et al., 2014; Ross et al., 2016). We recently showed that plasma level of psilocybin’s active metabolite psilocin is tightly coupled to both cerebral 5-HT2AR occupancy and the psychedelic experience (Madsen et al., 2019), and that baseline cerebral 5-HT2AR binding correlates with subjective effects of psilocybin (Stenbæk et al., 2020). These findings support that psilocin is a crucial pharmacological mediator of acute psilocybin effects and implicate the 5-HT2AR stimulation as mechanistically important. Considering the transformative potential of psilocybin as a transnosological treatment modality, directly linking psilocin and psilocybin-induced alterations in brain function can highlight critical drug-related mechanisms of psychedelic phenomenology and clinical outcome, yet has not been evaluated.

Acute psilocybin effects have been studied in four BOLD fMRI resting-state functional connectivity (RSFC) experiments: 1) 2 mg i.v. psilocybin in fifteen subjects (Carhart-Harris et al., 2012), 2) 0.2 mg/kg oral psilocybin in 23 subjects (Preller et al., 2020), 3) 0.14 mg/kg in 15 subjects (Barrett et al., 2020) and 4) 0.17 mg/kg in 30 subjects (Mason et al., 2020). Important findings from the first experiment include reduced RSFC within the DMN (Carhart-Harris et al., 2012), largely increased between-network RSFC (Roseman et al., 2014), and increased global RSFC (Tagliazucchi et al., 2016). The second experiment assessed BOLD fMRI scans acquired during the ascent phase and reported a reduction in global brain connectivity (GBC) in associative regions and increased GBC in sensory regions (Preller et al., 2020). The third study reported altered claustrum RSFC (Barrett et al., 2020), and the fourth reported decreased within-network RSFC and increased between-network RSFC (Mason et al., 2020). Notably, these studies did not assess plasma psilocin level (PPL) while measuring brain function nor investigate effects throughout the acute psychedelic experience following oral psilocybin intake. This knowledge is needed for delineating the neuropsychopharmacology of psilocybin and may inform the development of optimized treatment regimens.

In the present study, we sought to elucidate the extent to which network integrity and segregation map on to PPL and phenomenal experience alterations induced by a clinically relevant dose of oral psilocybin in healthy participants. To accomplish this, we collected simultaneous data of BOLD fMRI images, subjective drug intensity (SDI) ratings (i.e., real-time self-report psychedelic phenomenal experience changes in relation to psilocybin) (Madsen et al., 2019) and PPL (via blood samples) at multiple time points throughout the psilocybin psychedelic experience (5-6 hours). This scan strategy makes it possible to map brain function, phenomenal experience and PPL throughout psilocybin’s three pharmacological and experiential phases: ascent, peak and descent (Brown et al., 2017; Griffiths et al., 2011; Hasler et al., 2004, 1997; Lindenblatt et al., 1998; Madsen et al., 2019).

We hypothesized that PPL and SDI would correlate negatively with both DMN and average within-network RSFC, reflecting reduced network integrity, and positively with average-between network RSFC, reflecting reduced network segregation. To more closely identify potential network-specific effects, we also evaluated relations of RSFC within and between individual networks with PPL and SDI, respectively. Lastly, to complement the spatially constrained networks analysis, we evaluated spatially unconstrained voxel-level associations of PPL and SDI with local correlation (LCOR)(Deshpande et al., 2009), a connectivity measure reflecting local changes in integrity, and global correlation (GCOR)(Alfonso Nieto-Castanon, 2020), which reflects changes in whole-brain connectivity (WBC) and thus measures changes in global segregation.

## Experimental Procedures

### Participants

Fifteen healthy individuals (mean age ± SD 34.3 ± 9.8 years, six females) with no or limited experience with psychedelic substances participated in the study (**Table S1**). All participants were thoroughly prepared and supported before, during and after psilocybin by two psychologists.

### Procedures

Participants underwent MRI data acquisition (T1-weighed structural image and BOLD fMRI) prior to and after psilocybin administration (approximate dose: 0.2 mg/kg (n=4), 0.3 mg/kg (n=11), relative dose mean ± SD=0.26 ± 0.04 mg/kg, absolute dose mean ± SD=19 ± 3.5 mg). Participants were blind as to whether they would receive psilocybin (including dose) or a non-psychedelic drug. Post-drug rs-fMRI scan acquisition was performed approximately 40, 80, 130 and 300 minutes after psilocybin (see **Fig S1** for study design). Immediately after each 10-minute rs-fMRI scan acquisition, participants verbally rated the perceived subjective drug intensity (SDI) of the experience (“How intense is your experience right now?”, 0-10 Likert scale: 0 = “not at all intense”, 10 = “very intense”). Following the SDI rating, a blood sample was obtained from an intravenous catheter to quantify plasma level of free (unconjugated) psilocin. Further description of the intervention is available in **SI**. At the end of the day, participants completed self-report questionnaires to assess the overall psychedelic experience. More detailed information about participants and procedures is available in **SI**.

The data presented here stem from a larger study, which was approved by the ethics committee for the capital region of Copenhagen (journal identifier: H-16028698, amendments: 56023, 56967, 57974, 59673, 60437, 62255) and Danish Medicines Agency (EudraCT identifier: 2016-004000-61, amendments: 2017014166, 2017082837, 2018023295). Data was collected from October 2018 through February 2020.

### Psilocin plasma concentrations

Plasma psilocin analyses were performed using ultra performance liquid chromatography and tandem mass spectrometry as previously described (Madsen et al., 2019).

### Magnetic resonance imaging

Structural and functional MRI data were acquired on a 3T Siemens Prisma scanner (Siemens, Erlangen, Germany), using a 64-channel head coil. Image preprocessing was done in SPM12 (http://www.fil.ion.ucl.ac.uk/spm), and denoising and functional connectivity estimation was performed using CONN (Whitfield-Gabrieli and Nieto-Castanon, 2012) (see **SI** for further information).

## Data analysis

### PPL and SDI time course

The time course of PPL and SDI was visualized for each individual and for a group approximation (**Fig 1a and b**). The relation between PPL and SDI (**Fig 1c**) was modelled using an E_max_-model (constraints: 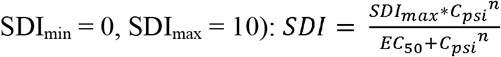, where “n” represents the Hill factor (Upton and Mould, 2014).

**Fig 1.**
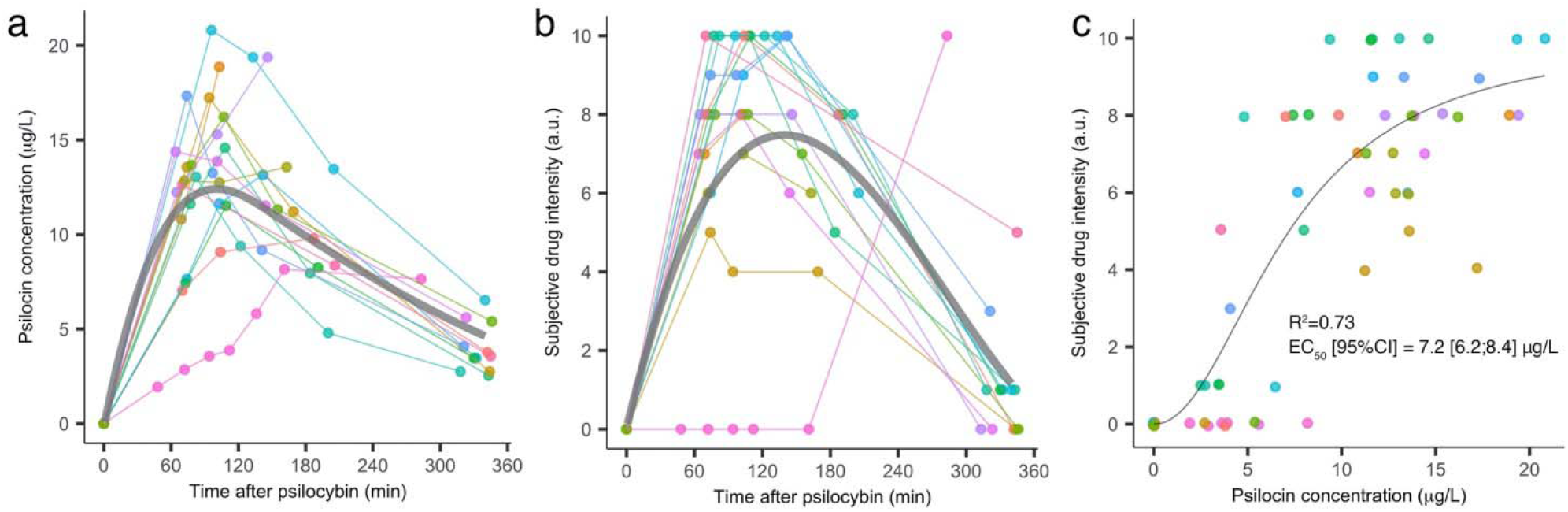
Plasma psilocin level and subjective drug intensity. Time course of plasma psilocin level (PPL) (a), and subjective drug intensity (SDI) (b). (c) Correlation between PPL and SDI. a.u., arbitrary units. Group time course of PPL and SDI approximated by spline fits is visualized in grey in (a) and (b). Datapoints are color-coded for each subject. One subject displayed a disjunction between PPL and SDI (color-coded in pink).

### Network analysis

Within-network and between FC was computed as the average of Fisher-transformed r-to-z values between regions within a given network or between a network pair for each rsfMRI acquisition. Linear mixed-effects regression analysis (Bates et al., 2015), which takes into account repeated observations for each subject, was used to model the association between FC and PPL and SDI, respectively (see **SI** for further information). Effects are reported in correlation coefficients (R) and unstandardized regression coefficients (β) with associated 95% confidence intervals (CI).

### Correction for multiple comparisons

Unless otherwise stated, the family-wise error rate (FWER) for all statistical tests was controlled at 5% using adjusted p-values (p_FWER_), where p_FWER_ below 0.05 was considered statistically significant (see **SI** for more details).

### Statistical analysis of GCOR and LCOR

Whole-brain GCOR and LCOR maps were included in separate linear mixed-effects models with PPL or SDI as independent variables, respectively. Based on the smoothness of the residuals, cluster sizes unlikely to occur by chance (p < 0.05) at a voxel-level threshold of p < 0.001 was estimated, using 3dFWHMx and 3dClustSim in AFNI version 16.2.01 (https://afni.nimh.nih.gov/). The cluster size threshold was calculated to be 560 voxels.

### Standard clinical dose response profile

A predicted response profile for a 25 mg clinical dose of psilocybin was constructed by spline fits of 0.3 mg/kg psilocin time course data from the present study, scaled to C_max_ of 18.7 μg/L, which is the approximate C_max_ of a 25 mg dose (Dahmane et al., 2020). Corresponding SDI and RSFC values were estimated using the predicted PPL curve and model parameters obtained in this study (**Fig 2** and **3**).

**Fig 2.**
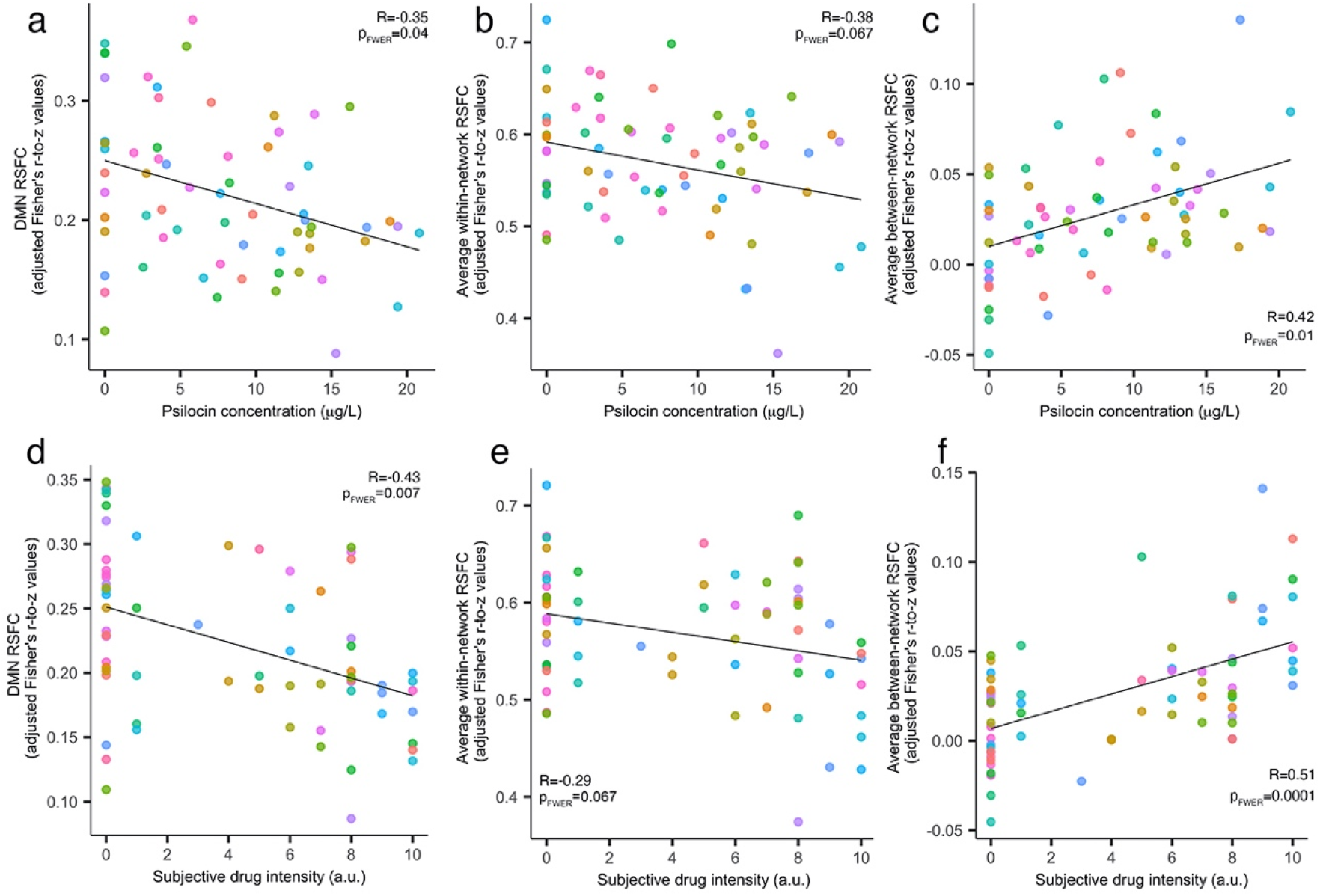
Network connectivity, plasma psilocin level and subjective drug intensity. Plasma psilocin level (PPL) was negatively associated with default mode network (DMN) resting-state functional connectivity RSFC (a) and average within-network RSFC (b), and positively associated with average between-network RSFC (c). Similar associations were observed for SDI (d, e, f). a.u., arbitrary units. adjusted: partial correlation for RSFC estimate on SDI or PPL. Datapoints are color-coded for each subject.

**Fig 3.**
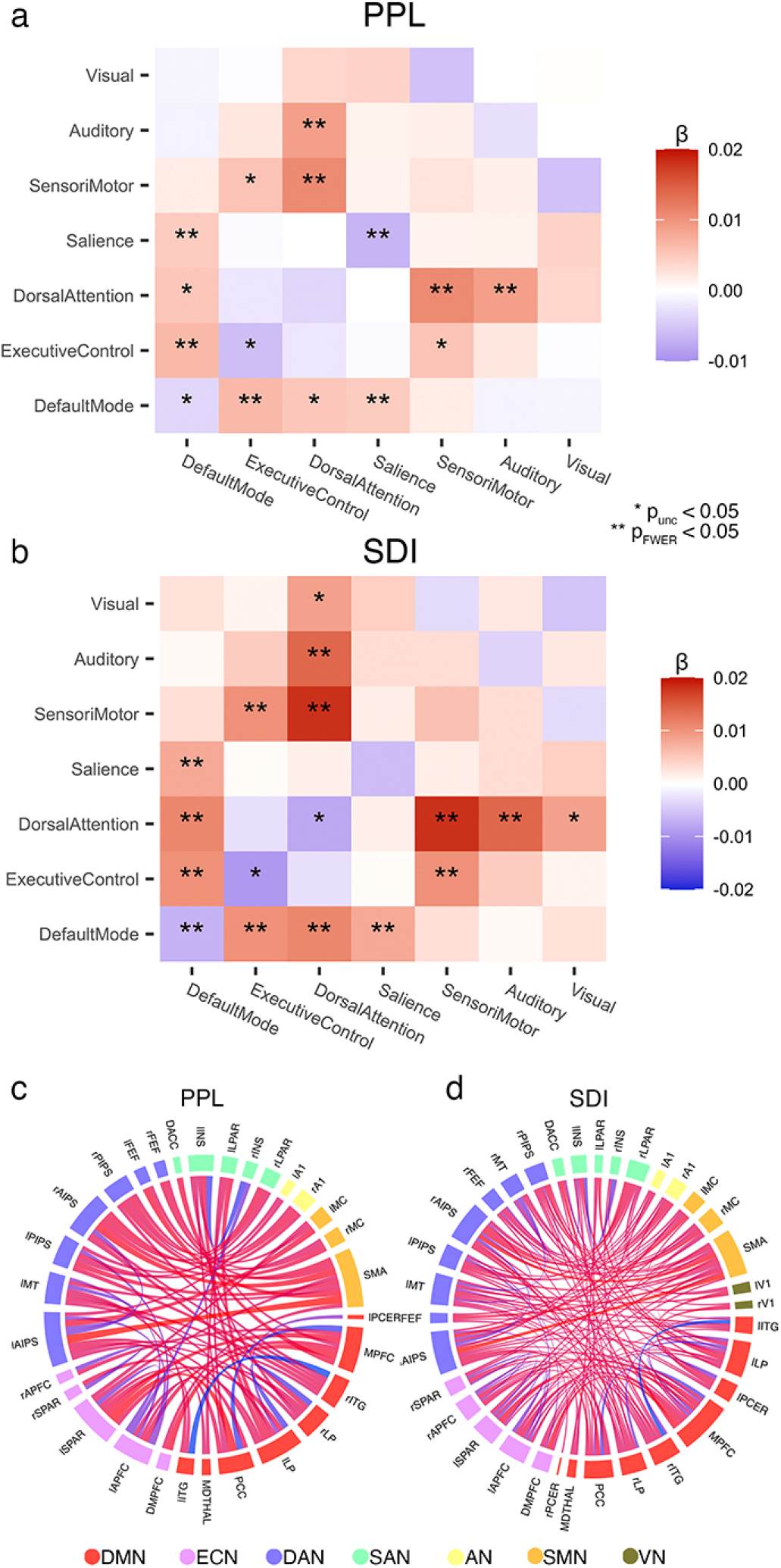
Results of individual networks analysis. Correlation matrix of the association between plasma psilocin level (PPL) and functional connectivity (FC) (a) and between subjective drug intensity (SDI) and FC (b). The diagonal elements represent association within a network while the off-diagonal elements represent the between-network FC of a pair of individual networks. * p_unc_ < 0.05, ** p_FWER_ < 0.05. A graphical representation is shown of the slope estimates (β) for the correlation of region-to-region FC with PPL (67 connections) (c) and with SDI (138 connections) (d), respectively; only estimates are shown for which q_FDR_ < 0.05. Red links signify a positive association; blue signifies a negative association. Line thickness is proportional to β estimate for each analysis (within each circle). Region names corresponding to abbreviations used in (c) and (d) are available in **SI Table 2**.

## Results

### Plasma psilocin level and subjective effects

In accordance with our previous findings (Madsen et al., 2019), PPL and SDI exhibited highly similar temporal trajectories and correlated positively (R^2^=0.73, EC_50_ [95%CI]=7.2 [6.2;8.4] μg/L, Hill factor=2.1) (**Fig 1**). One subject stood out by displaying slow pharmacokinetics and SDI insensitivity to psilocin at low PPL (**Fig 1**, color-coded in pink).

End-of-day retrospective self-report questionnaire assessment showed that the administered dose of psilocybin induced a profoundly altered state of consciousness, measured with the 11-Dimension Altered States of Consciousness questionnaire (11D-ASC) (Dittrich et al., 2006; Studerus et al., 2010), the revised Mystical Experiences Questionnaire (MEQ30) (Barrett et al., 2015) and the Ego-Dissolution Inventory (EDI) (Nour et al., 2016) (**Fig S2**). The area under curve (AUC) for SDI was statistically significantly positively correlated with average scores of 11D-ASC (R[95%CI]=0.59[0.14;0.85], β[95%CI]: 0.02[0.003;0.03], p_FWER_=0.028), total MEQ (R[95%CI]=0.57[0.08;0.84], β[95% CI]: 0.03[0.003;0.04], p_FWER_=0.028) and EDI (R[95%CI]=0.60[0.12;0.85], β[95% CI]: 0.02[0.006;0.06], p_FWER_=0.028) (**Fig S2**), consistent with our previous observation that SDI is sensitive to psilocybin-induced psychedelic phenomenology (Madsen et al., 2019).

### Associations of average network integrity and segregation with PPL and SDI

Seven resting-state networks were analyzed, comprising regions defined by 10 mm radius spheres about coordinates previously reported(Raichle, 2011) (36 regions-of-interest (ROIs), **Table S2**). Within- and between-network RSFC was estimated by averaging FC estimates between all pairs of relevant regions.

Although not statistically significant after correcting for multiple comparisons, a negative correlation was observed for average within-network RSFC (PPL: R[95%CI]=-0.28[-0.51;-0.01], β[95% CI]=-0.003[-0.006;-0.0002], p_FWER_=0.067; SDI: R[95%CI]=-0.29[-0.51;-0,02], β[95% CI]=-0.005[-0.009;-0.0005], p_FWER_=0.067) (**Fig 2**). PPL and SDI were statistically significantly positively correlated with average between-network RSFC (PPL: R[95%CI]=0.42[0.19;0.60], β[95%CI]=0.002[0.0008;0.004], p_FWER_=0.01; SDI: R = 0.59[0.37;0.74], β[95% CI]=0.005[0.003;0.007], p_FWER_=0.0001), reflecting reduced network segregation (**Fig 2**).

### Associations of integrity and segregation of individual networks with PPL and SDI

To better resolve the effects of PPL and SDI on the functional network architecture, we evaluated their associations with RSFC of individual networks and between network pairs (**Fig 3**). As hypothesized, both PPL and SDI were statistically significantly negatively correlated with DMN (PPL: R[95%CI]=-0.35[-0.61;-0.09], β[95% CI]: −0.004[-0.006;-0.00l], p_FWER_=0.04; SDI: R[95%CI]=-0.43[-0.65;-0.23], β[95% CI]=-0.007[-0.01;-0.003], p_FWER_=0.007), reflecting reduced network integrity (**Fig 2**). In addition to the DMN, integrity of the salience network (SAN) was also statistically significantly negatively correlated with PPL (**Fig 3a**; see **Table S3** for parameter estimates and statistics). Between-network analyses showed that as a function of PPL, segregation was statistically significantly reduced for 1) DMN with both SAN and executive control network (ECN, corresponding to the frontoparietal control network (Vincent et al., 2008; Yeo et al., 2011)), and 2) dorsal attention network (DAN) with the auditory network (AN) and sensorimotor network (SMN) (**Fig 3a; Table S3**). Segregation was statistically significantly reduced as a function of SDI for 1) DMN with SAN, ECN and DAN, 2) ECN with SMN, and 3) DAN with AN and SMN (**Fig 3b**; **Table S4**).

### Local correlation associations with PPL and SDI

LCOR measures FC between each voxel and its local neighborhood voxels. The LCOR analysis showed that PPL correlated negatively with LCOR in several regions belonging to DMN (medial prefrontal cortex (MPFC), posterior cingulate cortex (PCC) and precuneus), ECN (bilateral anterior prefrontal cortex (APFC) and left lateral parietal cortex (lLPC)), and visual cortex (occipital pole and lateral occipital cortex) (**Fig 4a**; **Table S5**). SDI was similarly statistically significantly negatively correlated with LCOR in regions belonging to DMN (PCC and precuneus), ECN (APFC and lLPC), visual cortex, and in bilateral temporal regions (**Fig 4a**; **Table S6**).

**Fig 4.**
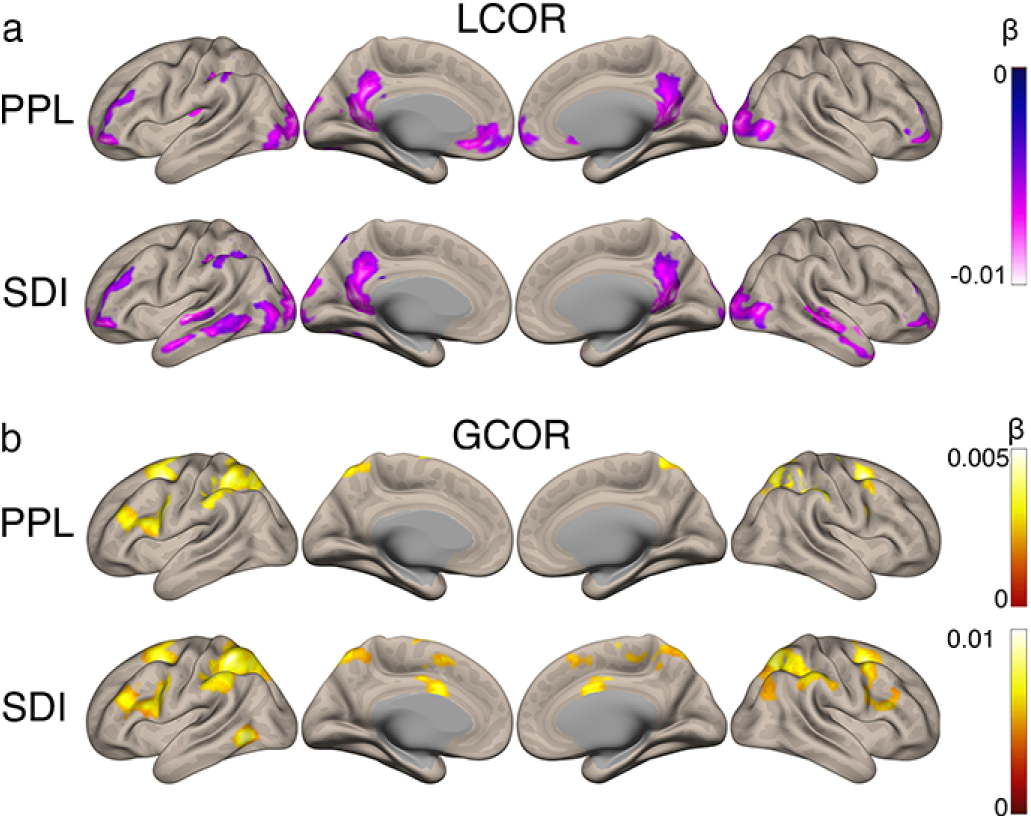
Results of the local correlation (LCOR) and global correlation (GCOR) analyses. Surface projection of areas for which LCOR (a), two top rows, and GCOR (b), two bottom rows, are significantly correlated with plasma psilocin level (PPL, upper rows) and subjective drug intensity (SDI, lower rows). LCOR associations were observed in visual cortex; in regions within default mode network hubs of medial prefrontal cortex (MPFC), posterior cingulate cortex (PCC) and temporal cortex; and in bilateral anterior prefrontal cortex (APFC), which is part of the ECN. GCOR correlated positively with SDI and PPL in clusters corresponding primarily to dorsal attention network (DAN) regions and to a lesser extent ECN. Color bars designate slope estimates (β) for the association. Only clusters larger than 560 voxels (voxel-level threshold p < 0.001) are shown.

### Global correlation associations with PPL and SDI

GCOR measures average FC between every voxel and the brain and provides a similar measure of WBC as used in previous psychedelic neuroimaging studies (Preller et al., 2020; Tagliazucchi et al., 2016). PPL showed statistically significant positive associations with GCOR in areas belonging to DAN and ECN (bilateral superior frontal gyri and bilateral intraparietal sulci) (**Fig 4b**; **Table S7**). SDI exhibited similar associations that also included a cluster in the dorsomedial cingulate gyrus, the supplementary motor area and thalamus (**Fig 4b and S3**; **Table S8**).

## Discussion

Here we evaluated how plasma psilocin and the psychedelic experience map onto RSFC measures of functional network integrity and segregation. As hypothesized, PPL correlated negatively with DMN network integrity. A similar negative correlation was observed for the SAN. The association with overall network integrity was similarly negative, albeit not statistically significant (p_FWER_ = 0.067). Similar associations were observed for SDI, implicating functional network integrity in phenomenal experience. PPL and SDI correlated negatively with LCOR in regions central to the DMN and ECN and in visual cortex—all regions with high 5-HT2AR levels (Ettrup et al., 2014). These findings suggest that network integrity is also impaired locally and are consistent with previous studies (Carhart-Harris et al., 2012; Mason et al., 2020; Muthukumaraswamy et al., 2013). Our results provide novel empirical evidence that psilocin shapes the integrity of DMN and SAN and implicate these neuropharmacological dynamics in the transformation of consciousness from the normal to the psychedelic state.

Consistent with our hypotheses, overall network segregation decreased as a function of PPL and SDI (**Fig 2**). This shows that functional networks increasingly synchronize as psilocin engages cerebral targets, including the 5-HT2AR (Madsen et al., 2019). This finding of reduced network segregation is consistent with previous observations of a functionally more connected brain in the psychedelic state induced by psilocybin (Carhart-Harris and Friston, 2019; Mason et al., 2020; Roseman et al., 2014) and lysergic acid diethylamide (LSD) (Robin L. Carhart-Harris et al., 2016; Müller et al., 2018, 2017; Preller et al., 2018; Tagliazucchi et al., 2016). Interestingly, however, effects of PPL and SDI on individual network integrity and segregation were observed only for some networks (**Fig 3**). Specifically, effects of PPL and SDI on individual network integrity appeared more pronounced for higher-level networks such as DMN and SAN compared to lower-level unimodal networks such as SMN and AN. Further, desegregating effects between individual networks matched the repeating spatial motif of brain networks previously described in the literature (Buckner and DiNicola, 2019; Margulies et al., 2016b). In this motif, the DMN is situated at one end, juxtaposed to heteromodal networks (ECN, DAN or SAN). These heteromodal networks are, in turn, situated next to unimodal networks such as auditory or visual networks, which reside at the other end. We found that as a function of PPL and SDI, segregation is reduced in a similar pattern for 1) DMN with DAN, ECN and SAN, and 2) DAN and ECN with unimodal sensory networks. Thus, although global network segregation is reduced, this effect appears more strongly driven by desegregation of adjacent functional networks.

The GCOR analysis showed increased WBC as a function of PPL and SDI for frontoparietal and midline regions within the DAN and ECN(Raichle, 2011; Yeo et al., 2011). Thus, the spatially unconstrained GCOR analysis and the spatially constrained networks analyses converge, showing desegregation among DMN, ECN, DAN and sensory networks as PPL rises and psychedelic phenomenology intensifies. The DMN is believed to contribute to selfhood/ego (Lebedev et al., 2015) and internal mentation (i.e., cognition dependent on constructed representations with minimal external stimuli (Buckner and DiNicola, 2019; Greicius et al., 2003; Raichle et al., 2001; Shulman et al., 1997)); the DAN is important in externally oriented attention (Corbetta and Shulman, 2002; Fox et al., 2005), and the ECN and SAN have been implicated as regulators of attention (Dixon et al., 2018; Menon and Uddin, 2010). Interestingly, subjective psychedelic experience includes unitive experiences and the perceived reduction in both sense of self and of the borders between self and the external world (Carhart-Harris et al., 2013). It is possible that psilocin-induced disruption of higher level networks, including the DMN, coupled with a “flowing together” of normally segregated functional streams related to internally directed and externally directed attention contributes to the psychedelic unitive experience, which may be therapeutically important for clinical outcomes of psilocybin therapy (Roseman et al., 2017).

Two other fMRI studies used WBC analyses, which yield measures similar to the GCOR analysis (Preller et al., 2020; Tagliazucchi et al., 2016). The first study showed increased WBC for frontoparietal networks, SAN and DMN, but not DAN, after i.v. psilocybin (Tagliazucchi et al., 2016). The second study reported only widespread decreased WBC when global-signal regression (GSR) was not used but increased WBC in sensory regions and reduced in associative and subcortical regions when GSR was used (Preller et al., 2020). Although Tagliazucchi and colleagues in the former study did not identify DAN as exhibiting increased WBC after psilocybin, our findings agree that other frontoparietal regions increase WBC after psilocybin. Findings from the latter WBC study by Preller and colleagues are neither in line with our findings nor the findings of Tagliazucchi and colleagues. Interstudy discrepancies may be influenced by differences related to psilocin pharmacokinetics (i.v. vs oral psilocybin formulation), BOLD signal denoising procedures (e.g., use of GSR) (Murphy and Fox, 2017), head motion or analytical method. Although future studies are warranted to more firmly establish psilocybin effects on voxel-level GBC, an important strength of our study resides in the direct linking of WBC with psilocin-induced changes in neurotransmission and phenomenal experience (as indexed by PPL and SDI).

The recently proposed ‘‘relaxed beliefs under psychedelics (REBUS) and the anarchic brain” theory hypothesizes that a core mechanism of 5-HT2AR agonist psychedelic compounds is impairment of the function of DMN and other high order networks (e.g., ECN), resulting in a less ordered (more entropic) and more desegregated functional architecture (Carhart-Harris and Friston, 2019). Importantly, we show that psilocin dose-dependently reduces network integrity and segregation (**Fig 2–4**), and our findings are thus compatible with neurobiological mechanisms proposed in the REBUS hypothesis.

Two recent studies showed reduced DMN integrity after challenge with the κ-opioid receptor agonist salvinorin A (psychoactive constituent of salvia divinorum) (Doss et al., 2020) and 3,4-methylenedioxymethamphetamine (MDMA) (Müller et al., 2021). DMN RSFC was also decreased after separate challenge with ketamine and midazolam (Adhikari et al., 2020). Thus, it is clear that reductions in RSFC within the DMN is not unique to 5-HT2AR agonist psychedelics. However, there is an overlap in the subjective experiences induced by MDMA (Studerus et al., 2010), salvia divinorum (González et al., 2006), psilocybin (Studerus et al., 2010), ketamine (Studerus et al., 2010) and LSD (Liechti et al., 2017), and it would be interesting to learn the extent to which changes in network integrity (including the DMN) and segregation map onto aspects of phenomenal experience induced by different psychoactive compounds.

Interestingly, the GCOR analysis also showed that thalamic GCOR correlated positively with SDI (**Fig S3**), compatible with previously observed substantive increases in thalamic RSFC after LSD (Müller et al., 2017). These findings are compatible with the thalamic gating theory of psychedelic drug action, which posits that thalamic filtering of external and internal information to the cortex is impaired by psilocin, resulting in information overload (Geyer and Vollenweider, 2008). Our observation of SDI-dependent increase in thalamic GCOR indicates that impaired thalamic gating contributes to psychedelic experiences in the context of psilocybin.

We replicate our previous finding that PPL and SDI display similar temporal trajectories and correlate positively (**Fig 1**) (Madsen et al., 2019). Notably, one participant stood out by exhibiting a “break-through” response style, i.e., delayed but rapid increase in SDI ratings (**Fig 1**, subject color-coded in pink). We speculate this may be explained at least partly by slow pharmacokinetics in this individual. Our observation that PPL and SDI correlate suggests that PPL is a an objective and quantitative determinant of overall subjective experience after psilocybin. Our results also support our previous observation that SDI correlates positively with retrospective questionnaire assessments of the psychedelic experience, including global ASC questionnaire score, EDI and MEQ30 (**Fig S2**). This indicates that SDI measures appraised integrated perceptual changes related to psychedelic phenomenology.

Consistent with previous psilocybin pharmacokinetics studies employing similar peroral doses (0.2-0.3 mg/kg) (Brown et al., 2017; Hasler et al., 1997; Lindenblatt et al., 1998), we observed considerable interindividual variability in maximum PPL (C_max_) and time to reach C_max_ (t_max_) (**Table S9**). Considering this pharmacological variability and the strong correlations of PPL with both phenomenal experience and FC changes, it is likely that interindividual differences in subjective effects and clinical response, which sometimes has been ascribed to psychological factors, are at least partly explained by interindividual differences in PPL (R. L. Carhart-Harris et al., 2016; Griffiths et al., 2016, 2011; Ross et al., 2016). Thus, our results indicate that future psilocybin studies would benefit from measuring PPL.

Our study is not without limitations. A larger sample size would have made it possible to detect more subtle effects. However, multiple measurements were done for each subject over time; such a strategy increases statistical power. A placebo condition could have helped identify potential expectancy effects on RSFC and SDI not tied to psilocybin. Nevertheless, participants were blind to the receipt of psilocybin and dose. Further, our study evaluated associations between RSFC, PPL and SDI, which are not necessarily inflated by an upward bias due to expectancy. It is difficult to imagine how placebo could mimic the tight associations with PPL across a similar time course and magnitude. Head motion correlated positively with PPL (**Fig S4**) but was generally of small magnitude, and quality control functional connectivity (QC-FC) plots indicated that noise was successfully removed in included, but not in excluded, datasets as part of the denoising procedure. Nevertheless, head motion effects need to be considered carefully. Hysteresis effects, i.e., different associations during ascent and descent phases, can confound our analysis strategy because we incorporated data points collected throughout the entire acute phase into a linear model. We note we have few data points to firmly exclude hysteresis effects but believe the problem is likely of smaller magnitude given the tight association of PPL and SDI and also the limited hysteresis effects of LSD (Dolder et al., 2015).

In conclusion, the present study evaluated real-time measures of PPL, alterations in phenomenal experience and FC measures of network integrity and segregation, covering ascent, peak and descent phases of medium to high and clinically relevant psilocybin doses in individuals with no or limited experience with psychedelic drugs. These findings demonstrate that psilocin critically shapes the time course and magnitude of changes in the cerebral functional architecture after psilocybin and implicate the expression of network integrity and segregation as important for consciousness and psychedelic experience.

## Contributors

PMF and GMK designed the study, wrote the protocol and contributed to data analysis, and interpretation. MKM designed the study, wrote the protocol, contributed to data collection, analysis, interpretation and wrote the first draft of the manuscript. MRMJ and AA assisted with data collection. DSS conceptualized the main psychological outcome of the study (SDI), and set up, translated and implemented the applied psychological measures and contributed with guiding at psilocybin interventions, participant preparation and integration. SA contributed with guiding at psilocybin interventions, and participant preparation and integration. KL and SSJ determined plasma psilocin levels. BO contributed to data analysis and interpretation. All authors contributed substantially to interpretation and discussion of the study results, critical review of the submitted manuscript, approve of the final version and agree to be both personally accountable for own contributions and willing to participate in resolving any question that may arise regarding the present paper.

## Acknowledgements

We gratefully acknowledge the work of MRI assistants, Agnete Dyssegaard and Arafat Nasser for biobank management, Oliver Overgaard-Hansen and Vibeke Dam for guiding at psilocybin interventions. We thank Lone Freyr, Gerda Thomsen, Svitlana Olsen, Peter Jensen and Dorthe Givard for technical/administrative assistance. We also acknowledge the BAFA laboratory, University of Chemistry and Technology and the National Institute of Mental Health (Prague, Czech Republic) for production of psilocybin and Glostrup Apotek (Glostrup, Denmark) for encapsulation.

