## Supplementary Materials for "Psilocybin-induced changes in brain network integrity and segregation correlate with plasma psilocin level and psychedelic experience"

DENMARK

**This document contains:**

Supplementary Materials and Methods

Figures S1 to S3

Tables S1 to S9

SI References

**Supplementary Materials and Methods**

**Psilocybin interventions.** Oral psilocybin (gelatin capsules each containing 3 mg pure psilocybin and glucose monohydrate as a filler) was administered shortly after the pre-drug scan with a glass of water. Participants were blind to whether they would receive psilocybin or a non-psychedelic drug and dose but were prepared for a potential experience of strong psychoactive effects by both the primary guide (psychologist) and a medical doctor. For the first subject, seven rounds of post-drug rs-fMRI data acquisitions were obtained. Although this subject did not report substantial distress due to the number of acquired scans, the acquisition plan was adjusted for all following participants (number of post-drug scans = 4) to provide participants a break from fMRI scan acquisition after the third post-drug rs-fMRI scan. Post-drug rs-fMRI data from the 300 min acquisition was missing for two subjects (one due to technical problems with the scanner, the other due to the participant feeling nauseous).

Two psychologists provided psychosocial support throughout the intervention day. At least one of the two psychologists met with participants the day after the intervention to allow for integration of the experience. Participants listened to music when in a room adjacent to the MRI scanner room from the time of psilocybin administration to the first post-drug acquisition and between the 130 and 300 min rs-fMRI acquisitions (i.e., during the break). No music was played in the context of rs-fMRI acquisition. After the 300-min scan session and when SDI was rated 1 or less, participants filled out three questionnaires retrospectively measuring aspects of the psychedelic experience: 11D-ASC [1,2], MEQ30 [3] and EDI[4].

Mental and somatic effects of psilocybin were as expected and no incidences of substantial discomfort or anxiety occurred. However, one participant experienced a vasovagal reaction app. two hours after psilocybin intake and experienced nausea and orthostatic hypotension. A similar occurrence was reported previously by another lab [5]. The participant in our study has previously taken psilocybin without such symptoms and indicated after this incident that he experienced a mild feeling of vertigo during a previous MRI scan conducted without any intervention. The participant did not express negative mental or somatic effects at the end of the intervention day, the day after, and one and two weeks after the intervention.

**Participants.** Volunteers were recruited from a list of volunteers who signed up to participate in a neuroimaging experiment investigating psilocybin. The data are part of a broader neuroimaging study; only procedures and results pertaining to psilocybin interventions are presented in the present paper. Prior to obtaining written informed consent, participants were informed about the study, side-effects and risks. After written informed consent was obtained, participants underwent a screening process, including screening for neurological illness or significant somatic illness, and a screening interview for present or previous psychiatric disorders using Mini-International Neuropsychiatric Interview, Danish translation version 6.0.0[6]. Exclusion criteria were: 1) present or previous primary psychiatric disease (DSM axis 1 or WHO ICD-10 diagnostic classifications) or in first-degree relatives; 2) present or previous neurological condition/disease, significant somatic condition/disease; 3) intake of drugs suspected to influence test results; 4) non-fluent Danish language skills; 5) vision or hearing impairment; 6) present or previous learning disability; 7) pregnancy; 8) breastfeeding; 9) MRI contraindications; 10) alcohol or drug abuse; 11) allergy to test drugs; 12) significant exposure to radiation within the past year; 13) intake of QT-prolonging medication or electrocardiogram (ECG) results indicative of heart disease; 14) blood donation less than three months before project participation; 15) bodyweight lower than 50 kg; 16) low plasma ferritin levels (< 12 μg/L). Self-reported history of hallucinogenic drug use was obtained and is presented in **Table S1**. Before the intervention day, participants were further prepared by the psychologist that would be the primary guide. Prior to fMRI data acquisition, a urine sample was obtained from each participant and subjected to a dip-stick test for common drugs of abuse (Rapid Response, BTNX Inc., Markham, Canada).

**MRI acquisition.** MRI data was obtained on a 3T Siemens Prisma scanner (Siemens, Erlangen, Germany), using a 64-channel head coil. BOLD fMRI data was obtained using a T2*-weighted gradient echo-planar imaging (EPI) sequence (TR = 2000 ms, TE = 30 ms, flip angle = 90^o^, in-plane matrix = 64x64 mm, in-plane resolution=3.6x3.6 mm, 32 slices (thickness = 3.0 mm, gap = 0.75 mm). Three hundred volumes were acquired for each BOLD fMRI data acquisition (10 min). Each participant was instructed to keep eyes closed, let the mind wander freely and not fall asleep. A high-resolution, T1-weighted 3D structural image was acquired at the pre-drug scan (inversion time = 900 ms, TE = 2.58 ms, TR = 1900 ms, flip angle = 9°, in-plane matrix = 256x256, resolution = 0.9x0.9 mm, 224 slices; slice thickness = 0.9 mm, no gap).

**fMRI data preprocessing.** Preprocessing of the BOLD fMRI data was performed using SPM12 (http://www.fil.ion.ucl.ac.uk/spm). The preprocessing steps included 1) realignment of functional images to first functional image, 2) co-registration of the T1-weighted structural image, 3) slice-timing correction, 4) unwarping, 5) normalization of images into Montreal Neurological Institute (MNI) space, 6) smoothing (8 mm Gaussian kernel), and 7) reslicing into a final voxel size of 2 x 2 x 2 mm.

**fMRI data denoising.** Previous neuroimaging studies report increased head motion after psychedelic drug intake [7–9], and head motion is potentially problematic as it may affect FC estimates and thus analytical results [10]. Drug-associated changes in respiratory or cardiac activity may likewise impact image quality and hence FC results [11]. Thus, it is important to clean the data from artifacts related to head motion and physiological noise. Head motion was limited by firmly fixing each subject’s head inside the head coil using foam pads. Post-acquisition denoising performed in CONN [12] included regression of noise sources at the level of data from each scan by including in the general linear model: 1) six motion parameters (three translation, three rotation) and their first-order derivatives, 2) the first five principal components and their first-order derivatives from separate principal components analyses of white matter and cerebrospinal fluid time series (aCompCor) [11,13], and 3) a regressor denoting images for which maximum voxel displacement exceeded 0.5 mm or for which global signal exceeded three standard deviations above the time series mean (spike regression) using ARTifact Detection Tools (ART, https://www.nitrc.org/projects/artifact_detect). Both spike regression [10,14,15] and aCompCor [10,11,13] improves BOLD fMRI data quality and has the benefit compared to GSR of not including gray matter signal, which can remove true neuronal signal [16–18]. The residuals represent the BOLD signal unexplained by the noise sources, i.e., the BOLD signal cleaned from noise. Functional datasets with less than 120 uncensored images (four minutes) were excluded from further analysis as previously described [10,15]. This procedure resulted in the exclusion of six datasets from the analysis, resulting in 68 included fMRI datasets.

Histograms of FC for a 1000-region adjacency matrix (QC-FC plots) computed before and after denoising were inspected as part of the denoising assessment [19]. The QC-FC plots for the omitted scans indicated that these scans indeed were tainted, even after denoising, supporting the exclusion of these datasets. The remaining QC-FC plots for the denoised BOLD data were unremarkable and suggested successful denoising.

**Functional connectivity estimation**

**Pairwise interregional analysis.** The network ROIs employed in the present analysis were defined by a 10 mm sphere about MNI coordinates previously reported [20] (see **Table S2)** for ROIs and coordinates used to delineate the networks). Estimation of FC was performed in CONN version 17.c (https://web.conn-toolbox.org/). The denoised BOLD signal time series for each ROI (percent of mean signal) was included in linear regression models for each 10-min rsfMRI acquisition, and the Fisher-transformed r-to-z correlation coefficients were calculated for all possible region pairs.

**LCOR analysis.** LCOR is a measure of local FC [21,22]. It is calculated for each voxel by computing a weighted-average of Fisher-transformed r-to-z-value correlation coefficients between each voxel time series and the time series of neighboring voxels, where the weighting is defined by an 8 mm full-width at half-maximum Gaussian kernel.

**GCOR analysis.** GCOR is implemented in CONN and is a FC method that gives a voxel-wise measure of GCOR [22]. Each whole-brain time-series was represented as a 64-component single value decomposition. Each component embody, in a stepdown fashion, the greatest amounts of functional connectivity [23]. GCOR was calculated as the average of Fisher-transformed r-to-z-value correlation coefficients for each voxel and the 64-component time-series.

**Data analysis**

**PPL and SDI time course.** The approximate average group time curves for PPL and SDI were visualized by spline fits. To regularize the spline for the PPL time curve, dummy data points were added at 500 minutes (PPL = 2 µg/L) and at 600 minutes (PPL = 1 µg/L). These dummy data points were not included in any analysis and were only used to constrain the group spline fit.

**Network analysis.** Average within-network RSFC was calculated as the average of all individual within-network RSFCs. Between-network RSFC was calculated as the average of all possible connections between regions in a pair of networks. Average between-network FC was calculated as the average of between-network RSFC estimates. Linear mixed-effects model regression analysis [24] was used to model the association between FC and PPL and SDI, respectively, and p-values were calculated using Satterthwaite’s method for approximating degrees of freedom [25]. The reported correlation coefficient R denotes the strength of the fixed effect (i.e., the relation of RSFC with PPL or SDI) when adjusting for random effects (i.e., the correlation with the partial residuals obtained by removing the subject-specific intercept from the outcome). Estimates of uncertainty (95% confidence intervals) of correlation coefficient R and slope β are not adjusted for multiple comparisons.

**Correction for multiple comparisons.** Unless otherwise stated, the family-wise error rate (FWER) for all statistical tests was controlled at 5% using adjusted p-values (p_FWER_), where p_FWER_ below 0.05 was considered statistically significant. Adjusted p-values for AUC SDI associations with psychedelic questionnaire outcomes were calculated using Hochberg’s method [26], and the Bonferroni-Holm method was used in all other cases. Hochberg’s method controls the type-I error rate with greater statistical power than Bonferroni-Holm but is valid only in certain situations (e.g., only positive or null correlations).

For RSFC analyses, statistical testing was first conducted for the associations of PPL and SDI was with average within-network, average between-network and DMN RSFC (number of tests = 6). Second, PPL and SDI associations were evaluated separately with all within- and between-network RSFC estimates (number of tests = 28). Third, associations of PPL and SDI with ROI-to-ROI RSFC were performed separately for PPL and SDI (number of tests = 630) and were thresholded using false-discovery-rate-corrected p-values (q_FDR_ < 0.05) [27].

**Supplementary Figures**

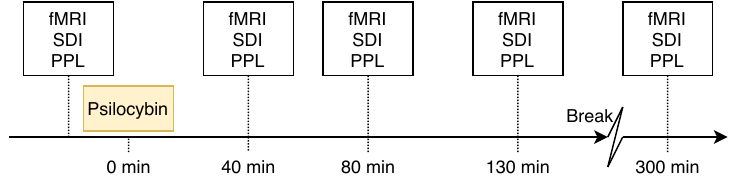

**Fig S1. Study design.** fMRI: functional magnetic resonance imaging, SDI: subjective drug intensity rating, PPL: plasma psilocin levels. After the third post-drug scan acquisition, participants had a break from fMRI acquisition before the fourth post-drug scan acquisition. Post-drug time designates anticipated beginning of scan.

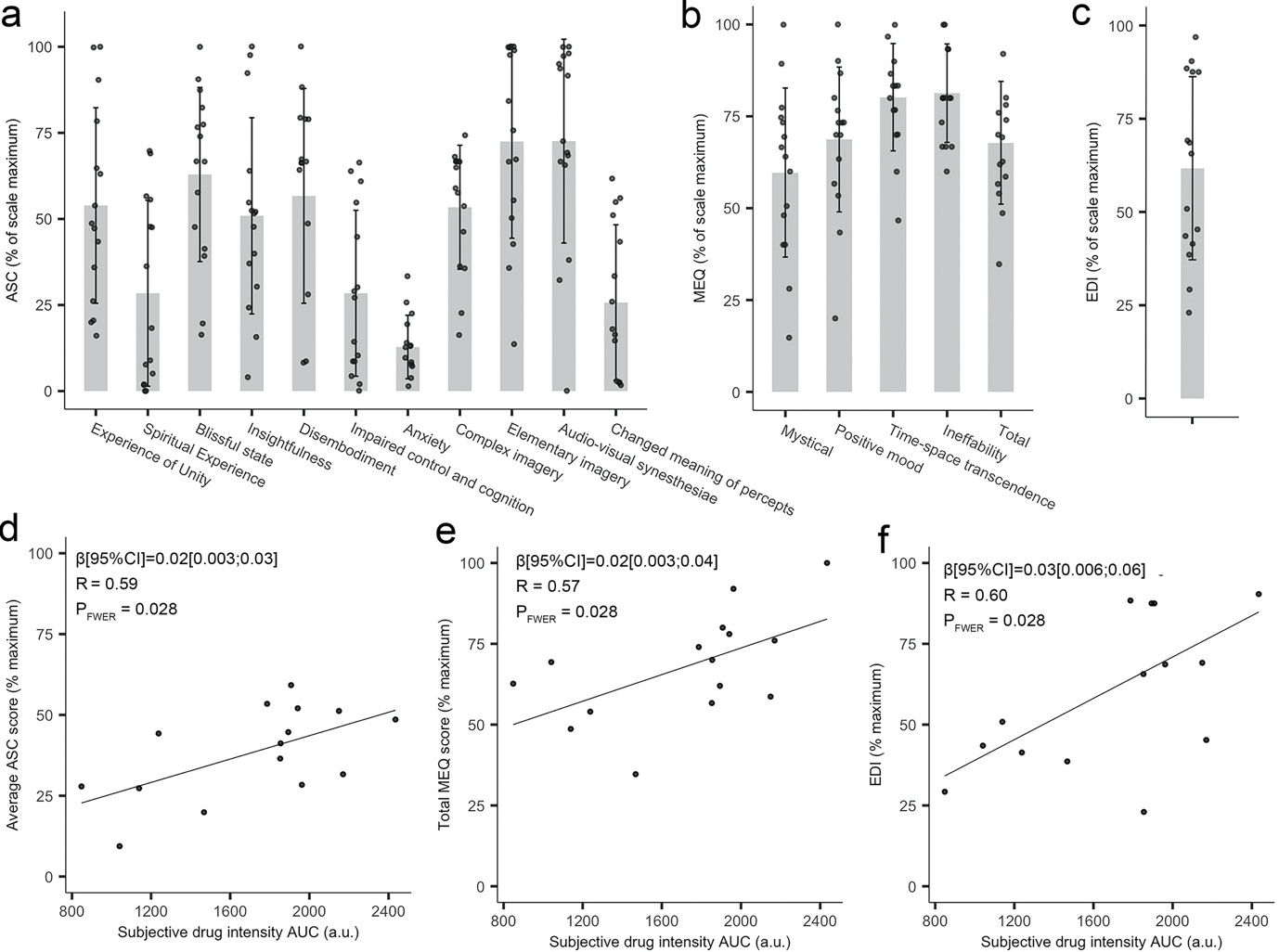

**Fig S2. Retrospective psychedelic questionnaire results and relation to subjective drug intensity.** Self-report responses for three questionnaires, measuring aspects of the psychedelic experience: (a) 11-Dimension Altered States of Consciousness (ASC), (b) Mystical Experiences Questionnaire (MEQ), (c) Ego-Dissolution Inventory (EDI). Correlation between the subjective drug intensity (SDI) area under curve (AUC) and ASC (d), MEQ30 (e) and EDI (f). a.u., arbitrary units.

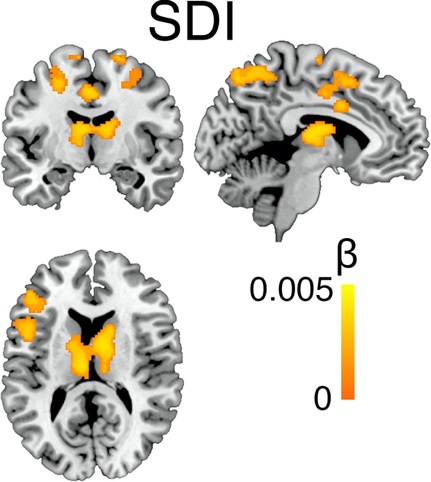

**Fig S3**. **Thalamus cluster identified in the analysis of GCOR association with SDI.** Color bar designate slope estimate (β) for the association. Only clusters larger than 560 voxels (voxel-level threshold p < 0.001) are shown. Image produced in MRIcron (https://www.nitrc.org/projects/mricron).

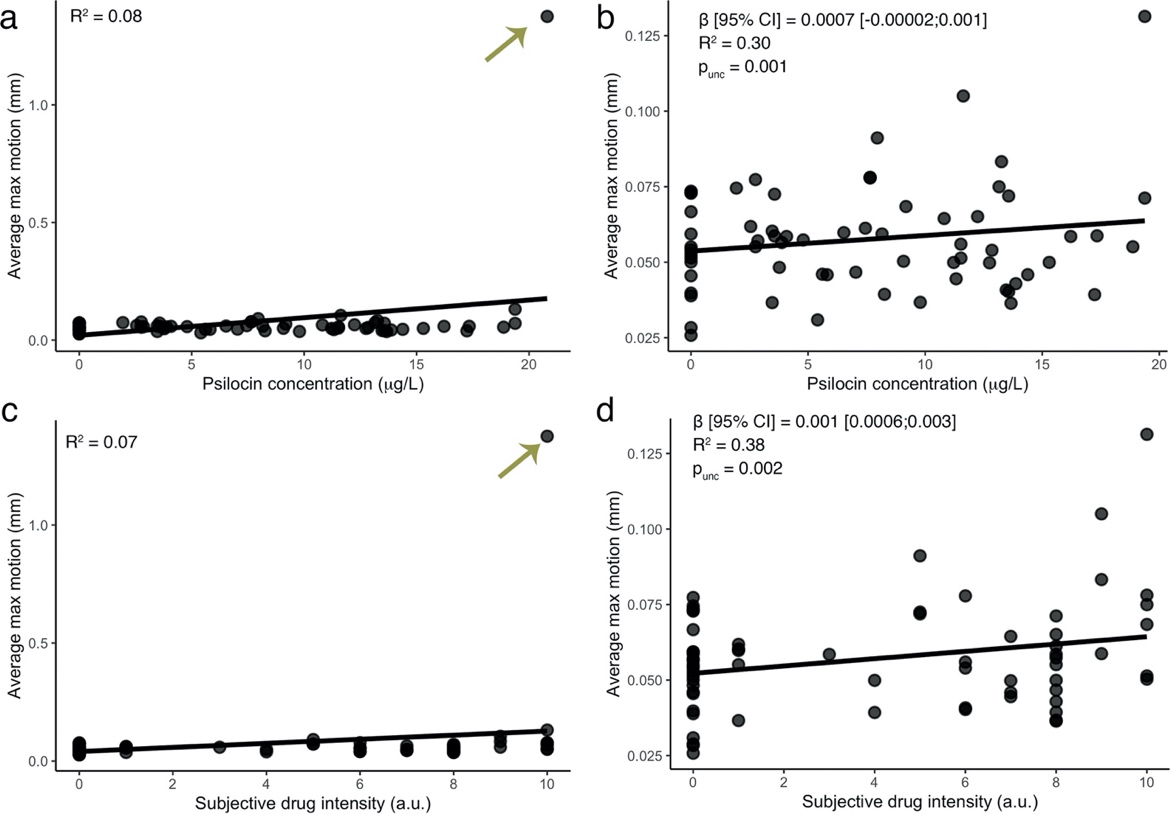

**Fig S4. Head motion, plasma psilocin levels (PPL) and subjective drug intensity (SDI).** Average head motion for each scan correlated positively PPL (b) and SDI (d) also when outlier (indicated by arrow) was removed (a) and (c). Previous neuroimaging studies report increased head motion after psychedelics.[7–9] Thus, we evaluated in the present sample the extent to which PPL and SDI correlate with head motion (the average of the greatest framewise voxel displacement for each BOLD fMRI scan acquisition (300 images per scan)). With the exception of one outlier (**SI Fig 2a** and **2c**), average maximum motion was low across all included datasets (mean ± SD: 0.08 ± 0.16 mm) in accordance with the procedure of firmly fixing each participant’s head inside the head coil using foam pads. Further, as expected, we observed a positive correlation of head motion with PPL and SDI (**SI Fig 2**). This finding suggests that it is important for neuroimaging studies investigating psilocybin to limit head motion and to employ good denoising procedures.

**Supplementary tables**

**Table S1. Participant history of hallucinogenic drug use.**

| **Subject** | **Life time** | **Preceding year** |
| --- | --- | --- |
| Subject 1 | No use | No use |
| Subject 2 | No use | No use |
| Subject 3 | No use | No use |
| Subject 4 | No use | No use |
| Subject 5 | No use | No use |
| Subject 6 | No use | No use |
| Subject 7 | *Psilocybin: 1* | No use |
| Subject 8 | Ketamine: 1 *Psilocybin: 1* | No use |
| Subject 9 | *Psilocybin: 1* | No use |
| Subject 10 | *Psilocybin: 1* | No use |
| Subject 11 | *Psilocybin: 1 Ayahuasca: 1* | No use |
| Subject 12 | *Psilocybin: 1* | No use |
| Subject 13 | No use | No use |
| Subject 14 | *Psilocybin: 9* | No use |
| Subject 15 | *Psilocybin: 1* | No use |

Self-reported history of use of hallucinogens. Serotonergic psychedelics italicized.

**Table S2. Region-of-interest-based networks delineation.**

|  |  |  | **MNI^a^ coordinate** |  |
| --- | --- | --- | --- | --- |
| **Network** | **Region** | **X** | **Y** | **Z** |
|  | Posterior cingulate cortex / precuneus | 0 | -51 | 27 |
|  | Medial prefrontal cortex | -1 | 54 | 27 |
|  | Left lateral parietal cortex | -46 | -66 | 30 |
| Default mode network | Right lateral parietal cortex | 49 | -63 | 33 |
|  | Left inferior temporal gyrus | -61 | -24 | -9 |
|  | Right inferior temporal gryus | 58 | -24 | -9 |
|  | Medial dorsal thalamus | 0 | -12 | 9 |
|  | Right posterior cerebellum | -25 | -81 | -33 |
|  | Left posterior cerebellum | 25 | -81 | 33 |
|  | Left frontal eye field | -29 | -9 | 54 |
|  | Right frontal eye field | 29 | -9 | 54 |
|  | Left posterior intraparietal sulcus | -26 | -66 | 48 |
| Dorsal attention network | Right posterior intraparietal sulcus | 26 | -66 | 48 |
|  | Left anterior intraparietal sulcus | -44 | -39 | 45 |
|  | Right anterior intraparietal sulcus | 41 | -39 | 45 |
|  | Left medial temporal | -50 | -66 | -6 |
|  | Right medial temporal | 53 | -63 | -6 |
|  | Dorsal medial prefrontal cortex | 0 | 24 | 46 |
|  | Left anterior prefrontal cortex | -44 | 45 | 0 |
| Executive control network | Right anterior prefrontal cortex | 44 | 45 | 0 |
|  | Left superior parietal cortex | -50 | -51 | 45 |
|  | Right superior parietal cortex | 50 | -51 | 45 |
|  | Dorsal anterior cingulate | 0 | 21 | 36 |
|  | Left anterior prefrontal cortex | -35 | 45 | 30 |
|  | Right anterior prefrontal cortex | 32 | 45 | 30 |
| Salience network | Left insula | -41 | 3 | 6 |
|  | Right insula | 41 | 3 | 6 |
|  | Left lateral parietal cortex | -62 | -45 | 30 |
|  | Right lateral parietal cortex | 62 | -45 | -40 |
|  | Left motor cortex | -39 | -26 | 41 |
| Sensorimotor network | Right motor cortex | 38 | -26 | 48 |
|  | Supplementary motor area | 0 | -21 | 48 |
| Visual network | Left V1 | -7 | 83 | 2 |
|  | Right V1 | 7 | 83 | 2 |
| Auditory network | Left A1 | -62 | -30 | 12 |
|  | Right A1 | 59 | -27 | 15 |

The table shows the center of mass coordinate for each 10 mm sphere used for delineation of each network, as described previously.[20] ^a^Montreal Neurological Institute coordinates.

**Table S3. Networks RSFC and plasma psilocin level (PPL).**

| **Network** | **Intercept** | **95% CI of intercept** | **ß-estimate** | **95% CI of ß** | **p_unc_** | **p_FWER_** |
| --- | --- | --- | --- | --- | --- | --- |
| AN | 0.8 | [0.68;0.92] | -0.0028 | [-0.011;0.0052] | 0.49 | 1 |
| DMN | 0.25 | [0.21;0.29] | -0.0036 | [-0.0065;-0.00081] | 0.014 | 0.04* |
| DAN | 0.44 | [0.39;0.48] | -0.0037 | [-0.0074;0.00035] | 0.059 | 1 |
| ECN | 0.42 | [0.36;0.49] | -0.0058 | [-0.01;-0.0013] | 0.014 | 0.294 |
| SAN | 0.42 | [0.37;0.47] | -0.0067 | [-0.01;-0.0029] | 0.00082 | 0.01968 |
| SMN | 0.45 | [0.36;0.54] | 0.0026 | [-0.0043;0.0088] | 0.42 | 1 |
| VN | 1.4 | [1.2;1.5] | 0.00019 | [-0.012;0.013] | 0.98 | 1 |
| DMN-AN | -0.09 | [-0.15;-0.029] | -0.0012 | [-0.0053;0.0028] | 0.56 | 1 |
| DMN-DAN | -0.15 | [-0.19;-0.1] | 0.0052 | [0.0016;0.0086] | 0.0032 | 0.0704 |
| DMN-ECN | 0.0022 | [-0.045;0.048] | 0.0065 | [0.0034;0.0095] | 5.70*10^-5^ | 0.001539 |
| DMN-SAN | -0.097 | [-0.14;-0.054] | 0.0048 | [0.0024;0.007] | 0.00013 | 0.00338 |
| DMN-SMN | -0.042 | [-0.077;-0.0072] | 0.0018 | [-0.0014;0.0052] | 0.27 | 1 |
| DMN-VN | 0.0062 | [-0.047;0.059] | -0.0011 | [-0.0056;0.0033] | 0.64 | 1 |
| DAN-AN | 0.081 | [0.018;0.14] | 0.0089 | [0.004;0.014] | 0.00066 | 0.0165 |
| DAN-ECN | 0.079 | [0.029;0.13] | -0.0022 | [-0.0061;0.0015] | 0.24 | 1 |
| DAN-SAN | 0.049 | [0.0045;0.092] | 2.30E-05 | [-0.0032;0.0032] | 0.99 | 1 |
| DAN-SMN | 0.093 | [0.041;0.14] | 0.011 | [0.0057;0.015] | 1.70*10^-5^ | 0.000476 |
| DAN-VN | 0.017 | [-0.049;0.085] | 0.0037 | [-0.0018;0.0094] | 0.19 | 1 |
| ECN-AN | -0.078 | [-0.13;-0.029] | 0.0023 | [-0.0019;0.0065] | 0.28 | 1 |
| ECN-SAN | 0.1 | [0.052;0.15] | -0.00056 | [-0.0037;0.0026] | 0.73 | 1 |
| ECN-SMN | -0.1 | [-0.13;-0.067] | 0.0055 | [0.0021;0.009] | 0.0025 | 0.0575 |
| ECN-VN | -0.047 | [-0.1;0.0097] | -0.00036 | [-0.0051;0.0044] | 0.88 | 1 |
| SAN-AN | 0.21 | [0.15;0.27] | 0.0014 | [-0.0038;0.0062] | 0.56 | 1 |
| SAN-SMN | 0.018 | [-0.027;0.061] | 0.0012 | [-0.0026;0.0049] | 0.53 | 1 |
| SAN-VN | -0.085 | [-0.13;-0.041] | 0.004 | [-0.00013;0.0084] | 0.063 | 1 |
| SMN-AN | 0.17 | [0.11;0.23] | 0.0016 | [-0.0043;0.007] | 0.56 | 1 |
| SMN-VN | 0.056 | [-0.019;0.13] | -0.0056 | [-0.012;0.00087] | 0.094 | 1 |
| VN-AN | 0.021 | [-0.067;0.11] | 8.70E-05 | [-0.0066;0.007] | 0.98 | 1 |

Result of linear mixed-effects model analysis of the association between PPL and functional connectivity within and between networks (networks separated by “-”). AN: auditory network; DMN: default mode network; DAN: dorsal attention network; ECN: executive control network; SAN: salience network; SMN: sensorimotor network; VN: visual network. FC: functional connectivity. The intercept represents the model-estimated FC at pre-drug and thus gives a measure of FC in the unstimulated state. 95% CI reflects confidence interval not adjusted for multiple comparisons. P_unc_ denotes the p-value unadjusted for multiple comparisons, and p_FWER_ is adjusted using Bonferroni-Holm. *Hypothesized *a priori* to correlate with PPL.

**Table S4. Networks RSFC and subjective drug intensity (SDI).**

| **Network** | **Intercept** | **95% CI of intercept** | **ß-estimate** | **95%CI of ß** | **p_unc_** | **p_FWER_** |
| --- | --- | --- | --- | --- | --- | --- |
| AN | 0.79 | [0.68;0.91] | -0.0039 | [-0.016;0.0083] | 0.53 | 1 |
| DMN | 0.25 | [0.22;0.28] | -0.0069 | [-0.011;-0.0029] | 0.0014 | 0.007* |
| DAN | 0.44 | [0.41;0.48] | -0.0078 | [-0.014;-0.0018] | 0.011 | 0.22 |
| ECN | 0.41 | [0.35;0.48] | -0.0093 | [-0.016;-0.0026] | 0.008 | 0.168 |
| SAN | 0.39 | [0.34;0.44] | -0.006 | [-0.012;0.00024] | 0.06 | 1 |
| SMN | 0.44 | [0.36;0.53] | 0.0058 | [-0.0041;0.015] | 0.23 | 1 |
| VN | 1.4 | [1.3;1.5] | -0.0053 | [-0.025;0.014] | 0.59 | 1 |
| DMN-AN | -0.1 | [-0.16;-0.041] | 0.00073 | [-0.0053;0.0068] | 0.81 | 1 |
| DMN-DAN | -0.16 | [-0.2;-0.12] | 0.011 | [0.0061;0.016] | 4.00*10^-5^ | 0.001 |
| DMN-ECN | 0.0067 | [-0.035;0.048] | 0.01 | [0.0059;0.014] | 1.40*10^-5^ | 0.000378 |
| DMN-SAN | -0.094 | [-0.14;-0.054] | 0.0079 | [0.0046;0.011] | 1.40*10^-5^ | 0.000378 |
| DMN-SMN | -0.041 | [-0.074;-0.0084] | 0.0029 | [-0.0019;0.008] | 0.24 | 1 |
| DMN-VN | -0.014 | [-0.065;0.037] | 0.0027 | [-0.0041;0.0093] | 0.43 | 1 |
| DAN-AN | 0.086 | [0.024;0.15] | 0.014 | [0.0065;0.021] | 0.00041 | 0.00943 |
| DAN-ECN | 0.075 | [0.026;0.12] | -0.0029 | [-0.0084;0.0028] | 0.32 | 1 |
| DAN-SAN | 0.041 | [-7.2e-05;0.082] | 0.0016 | [-0.0032;0.0063] | 0.51 | 1 |
| DAN-SMN | 0.096 | [0.049;0.14] | 0.018 | [0.011;0.025] | 2.50*10^-6^ | 7.00*10^-5^ |
| DAN-VN | 0.0096 | [-0.051;0.071] | 0.0087 | [0.00041;0.017] | 0.045 | 0.855 |
| ECN-AN | -0.08 | [-0.13;-0.035] | 0.0047 | [-0.0017;0.011] | 0.15 | 1 |
| ECN-SAN | 0.092 | [0.046;0.14] | 0.00042 | [-0.0043;0.0051] | 0.86 | 1 |
| ECN-SMN | -0.1 | [-0.13;-0.072] | 0.01 | [0.005;0.015] | 0.00026 | 0.00624 |
| ECN-VN | -0.055 | [-0.11;-0.0011] | 0.0011 | [-0.0061;0.0083] | 0.76 | 1 |
| SAN-AN | 0.21 | [0.16;0.27] | 0.0031 | [-0.0044;0.01] | 0.4 | 1 |
| SAN-SMN | 0.022 | [-0.019;0.063] | 0.0017 | [-0.0041;0.0076] | 0.57 | 1 |
| SAN-VN | -0.072 | [-0.11;-0.03] | 0.0044 | [-0.0021;0.011] | 0.19 | 1 |
| SMN-AN | 0.17 | [0.12;0.23] | 0.0031 | [-0.0052;0.011] | 0.45 | 1 |
| SMN-VN | 0.027 | [-0.044;0.1] | -0.0033 | [-0.013;0.0069] | 0.52 | 1 |
| VN-AN | 0.014 | [-0.07;0.1] | 0.0022 | [-0.0081;0.013] | 0.68 | 1 |

Result of linear mixed-effects model analysis of the association between SDI and functional connectivity within and between networks (networks separated by “-”). AN: auditory network; DMN: default mode network; DAN: dorsal attention network; ECN: executive control network; SAN: salience network; SMN: sensorimotor network; VN: visual network. FC: functional connectivity. The intercept represents the model-estimated FC at pre-drug and thus gives a measure of FC in the unstimulated state. 95% CI reflects confidence interval not adjusted for multiple comparisons. P_unc_ denotes the p-value unadjusted for multiple comparisons, and p_FWER_ is adjusted using Bonferroni-Holm. *Hypothesized *a priori* to correlate with SDI.

**Table S5. Results of analysis of local correlation (LCOR) and plasma psilocin level (PPL).**

| **Cluster** | **Regions** | **Size (voxels)** | **Coordinate of peak voxel** |
| --- | --- | --- | --- |
| 1  Negative association | Left middle occipital | 1325 | (-36,-82,-20) |
|  | Left superior occipital |  |  |
|  | Left calcarine |  |  |
|  | Left inferior occipital |  |  |
|  | Left fusiform |  |  |
| 2  Negative association | Left middle frontal, orbital part | 947 | (-8,30-10) |
|  | Left anterior cingulate |  |  |
|  | Right middle frontal, orbital part |  |  |
| 3  Negative association | Left middle frontal | 701 | (44,50,-8) |
|  | Right middle frontal |  |  |
|  | Right middle frontal, orbital part |  |  |
|  | Right inferior frontal, orbital part |  |  |
|  | Right inferior frontal, triangular part |  |  |
| 4  Negative association | Left middle frontal | 1079 | (-38,52,-8) |
|  | Left middle frontal, orbital part |  |  |
|  | Left inferior frontal, triangular part |  |  |
| 5  Negative association | Right middle occipital | 1443 | (20,-98,6) |
|  | Right superior occipital |  |  |
|  | Right inferior occipital |  |  |
| 6  Negative association | Right precuneus | 3053 | (14,-50,12) |
|  | Left precuneus |  |  |
|  | Left calcarine |  |  |
|  | Right calcarine |  |  |
|  | Left posterior cingulum |  |  |
|  | Left middle cingulum |  |  |
|  | Right posterior cingulum |  |  |
|  | Rihgt middle cingulum |  |  |

The table shows clusters within which there was a statistically significant correlation between local correlation (LCOR) and PPL. Coordinate in Montreal Neurological Institute (MNI) space. Cluster size threshold was 560 voxels. Only negative associations were observed.

**Table S6. Results of analysis of local correlation** (**LCOR) and subjective drug intensity (SDI).**

| **Cluster** | **Regions** | **Size (voxels)** | **Peak voxel coordinate** |
| --- | --- | --- | --- |
| 1  Negative association | Right middel temporal | 1045 | (52,-22,-8) |
|  | Right middle temporal pole |  |  |
|  | Right superior temporal |  |  |
| 2  Negative association | Left middel temporal | 3495 | (-64,-14,-6) |
|  | Left middle occipital |  |  |
|  | Left inferior temporal |  |  |
|  | Left superior occipital |  |  |
|  | Left inferior occipital |  |  |
|  | Left cuneus |  |  |
|  | Left calcarine |  |  |
| 3  Negative association | Right middle frontal | 1034 | (26,60,-8) |
|  | Right middle frontal, orbital part |  |  |
|  | Right inferior frontal, orbital part |  |  |
| 4  Negative association | Left middle frontal | 1391 | (-36,54,-8) |
|  | Left middle frontoorbital |  |  |
|  | Left inferior frontal triangular part |  |  |
|  | Left inferior frontal orbital part |  |  |
| 5  Negative association | Right middle occipital | 1380 | (48,-74,2) |
|  | Right superior occipital |  |  |
|  | Right inferior occipital |  |  |
|  | Right middle temporal |  |  |
|  | Right inferior temporal |  |  |
| 6  Negative association | Right precuneus | 3077 | (16,-52,14) |
|  | Left precuneus |  |  |
|  | Left posterior cingulate |  |  |
|  | Right posterior cingulate |  |  |
|  | Left middle cingulate |  |  |
|  | Right middle cingulate |  |  |
|  | Right calcarine |  |  |
|  | Left calcarine |  |  |
| 7  Negative association | Left inferior parietal | 1973 | (-38,-32,38) |
|  | Left superior parietal |  |  |
|  | Left middle occipital |  |  |
|  | Right precuneus |  |  |
|  | Left precuneus |  |  |
|  | Right superior parietal |  |  |

The table shows clusters within which there was a statistically significant correlation for LCOR with SDI. Coordinate in Montreal Neurological Institute (MNI) space. Cluster size threshold was 560 voxels. Only negative associations were observed.

**Table S7. Results of analysis of global correlation** (**GCOR) and plasma psilocin level (PPL).**

| **Cluster** | **Regions** | **Size (voxels)** | **Peak voxel coordinate** |
| --- | --- | --- | --- |
| 1  Positive association | Left inferior frontal, triangular part | 1372 | (-52,6,34) |
|  | Left inferior frontal, opercular part |  |  |
|  | Left middle frontal |  |  |
| 2  Positive association | Right middle frontal | 857 | (24,4,50) |
|  | Right superior frontal |  |  |
|  | Right middle cingulate |  |  |
|  | Right precentral |  |  |
| 3  Positive association | Left inferior parietal | 2616 | (-42,44,54) |
|  | Left superior parietal |  |  |
|  | Left precuneus |  |  |
| 4  Positive association | Right superior parietal | 907 | (20,-66,56) |
|  | Right precuneus |  |  |
|  | Right post central |  |  |
|  | Right inferior parietal |  |  |
| 5  Positive association | Right inferior parietal | 690 | (46,-38,52) |
|  | Right supramarginal |  |  |
|  | Right post-central |  |  |
| 6  Positive association | Left superior frontal | 757 | (-26,2,54) |
|  | Left middle frontal |  |  |
|  | Left pre-central |  |  |

The table shows clusters within which there was a statistically significant correlation between GCOR PPL. Coordinate in Montreal Neurological Institute (MNI) space. Only positive associations were observed. Cluster size threshold was 560 voxels.

**Table S8. Results of analysis of global correlation** **(GCOR) and subjective drug intensity (SDI).**

| **Cluster** | **Regions** | **Size (voxels)** | **Peak voxel coordinate** |
| --- | --- | --- | --- |
| 1  Positive association | Left inferior temporal | 563 | (-52,-60,-6) |
|  | Left middle temporal |  |  |
|  | Left inferior occipital |  |  |
|  | Left middle occipital |  |  |
| 2  Positive association | Left thalamus | 791 | (10,-2,14) |
|  | Right thalamus |  |  |
|  | Left caudate |  |  |
|  | Right caudate |  |  |
| 3  Positive association | Left inferior frontal, triangular part | 1794 | (-50,6,30) |
|  | Left precentral |  |  |
|  | Left inferior frontal, opercular part |  |  |
|  | Left middle frontal |  |  |
| 4  Positive association | Right middle frontal | 1977 | (24,4,50) |
|  | Right inferior frontal, opercular part |  |  |
|  | Right superior frontal |  |  |
|  | Right precentral |  |  |
|  | Right inferior frontal, triangular part |  |  |
| 5  Positive association | Left inferior parietal | 7862 | (-42,-46,56) |
|  | Right inferior parietal |  |  |
|  | Left superior parietal |  |  |
|  | Right superior parietal |  |  |
|  | Left precuneus |  |  |
|  | Right precuneus |  |  |
|  | Left supramarginal |  |  |
|  | Right superior occupital |  |  |
|  | Left superior occipital |  |  |
|  | Right middle occipital |  |  |
|  | Left middle occipital |  |  |
|  | Left post central |  |  |
|  | Right post central |  |  |
|  | Right middle cingulum |  |  |
|  | Left middle cingulum |  |  |
| 6  Positive association | Left sup. motor area | 778 | (2,6,30) |
|  | Left middle cingulate |  |  |
|  | Right sup. motor area |  |  |
|  | Right middle cingulate |  |  |
|  | Left anterior cingulate |  |  |
|  | Right anteriate cingulate |  |  |
| 7  Positive association | Left superior frontal | 929 | (-26,4,50) |
|  | Left middle frontal |  |  |

The table shows clusters within which there was a statistically significant correlation between GCOR and SDI. Coordinate in Montreal Neurological Institute (MNI) space. Only positive associations were observed. Cluster size threshold was 560 voxels.

**Table S9. Plasma psilocin level (PPL).**

| **Dose (mg/kg)** | **n** | **C_max_ median [range] (µg/L)** | **C_max_ mean** ± **SD(µg/L)** | **t_max_ median [range] (min)** | **t_max_ mean** ± **SD (min)** |
| --- | --- | --- | --- | --- | --- |
| 0.2 | 4 | 13.8 [8.2-17.3] | 13.3 ± 3.8 | 108 [64-161] | 110 ± 48 |
| 0.3 | 11 | 14.6 [9.8-20.9] | 15.2 ± 3.5 | 107 [70;187] | 115 ± 36 |

C_max_: maximum plasma psilocin concentration. t_max_: time to C_max_. PPL measured after each fMRI acquisition.
